# Localisation of Apicomplexa motor Myosin A with axial nanometric precision using graphene energy transfer

**DOI:** 10.1101/2025.07.30.667387

**Authors:** Giovanni Ferrari, Yuan Song, Michael Hensel, Olympia Ekaterini Psathaki, Doan Thuy Nguyen Nguyen, Simon Gras, Markus Meissner, Philip Tinnefeld, Javier Periz

## Abstract

Migration in Apicomplexan parasites, a phylum that includes Plasmodium spp. responsible for malaria and the zoonotic Toxoplasma, is explained by the linear migration motor model. This model predicts that Myosin A is confined beneath the plasma membrane and above the inner membrane complex (IMC), a membranous barrier separating the cell membrane and the cytoplasm. Cumulative data, using mutant cell lines and biochemical data support this model. Paradoxically, a proof of the precise localization of Myosin A is still lacking due to limitations in resolving the motor with nanometric precision within the space comprising the membrane and the IMC. Here, we implement graphene energy transfer (GET), a novel axial nanometric ruler with a resolution of ∼1 nm, to determine the relative axial position of Myosin A and IMC1 (a reference protein defining IMC position). Using GET, we estimate the IMC dimensions with precision matching electron microscopy (EM) data and the added advantage of identifying specific bauplan proteins. We complement these measurements with 2D STED microscopy in MIRA confiners, uExMIC, and cryo-immunolabeling. Our data present the first direct localization of Myosin A populations with nanometric resolution in the third dimension, which is compatible with the linear motor model.

**One sentence summary:** Using the novel GET method, this study presents the first direct localization of Myosin A populations in *Toxoplasma gondii* with 1-4 nm axial precision, supporting the linear model.

## Introduction

Apicomplexa is a large phylum of parasitic protists belonging to the Alveolata group and includes major pathogens such as *Plasmodium spp* the causative agent of malaria, or *Toxoplasma gondii* responsible for zoonotic toxoplasmosis. Apicomplexa have evolved a conserved migration mechanism, powered by actin and myosin motors to find and invade host cells ^1^.

A cumulative amount of data, lead to the identification of critical factors involved in gliding motility. The glideosome, a multi-protein complex consisting of MyoA, MLC1, GAP45, GAP50 and GAPM proteins is predicted to power motility. The linear motor model, postulates the glideosome is anchored ^2 3^ to the inner membrane complex (IMC) a structure that consists of membranous cisternae and structural components located 30-40 nm beneath the PM ^4^(Fig. 1a).

**Figure 1.**
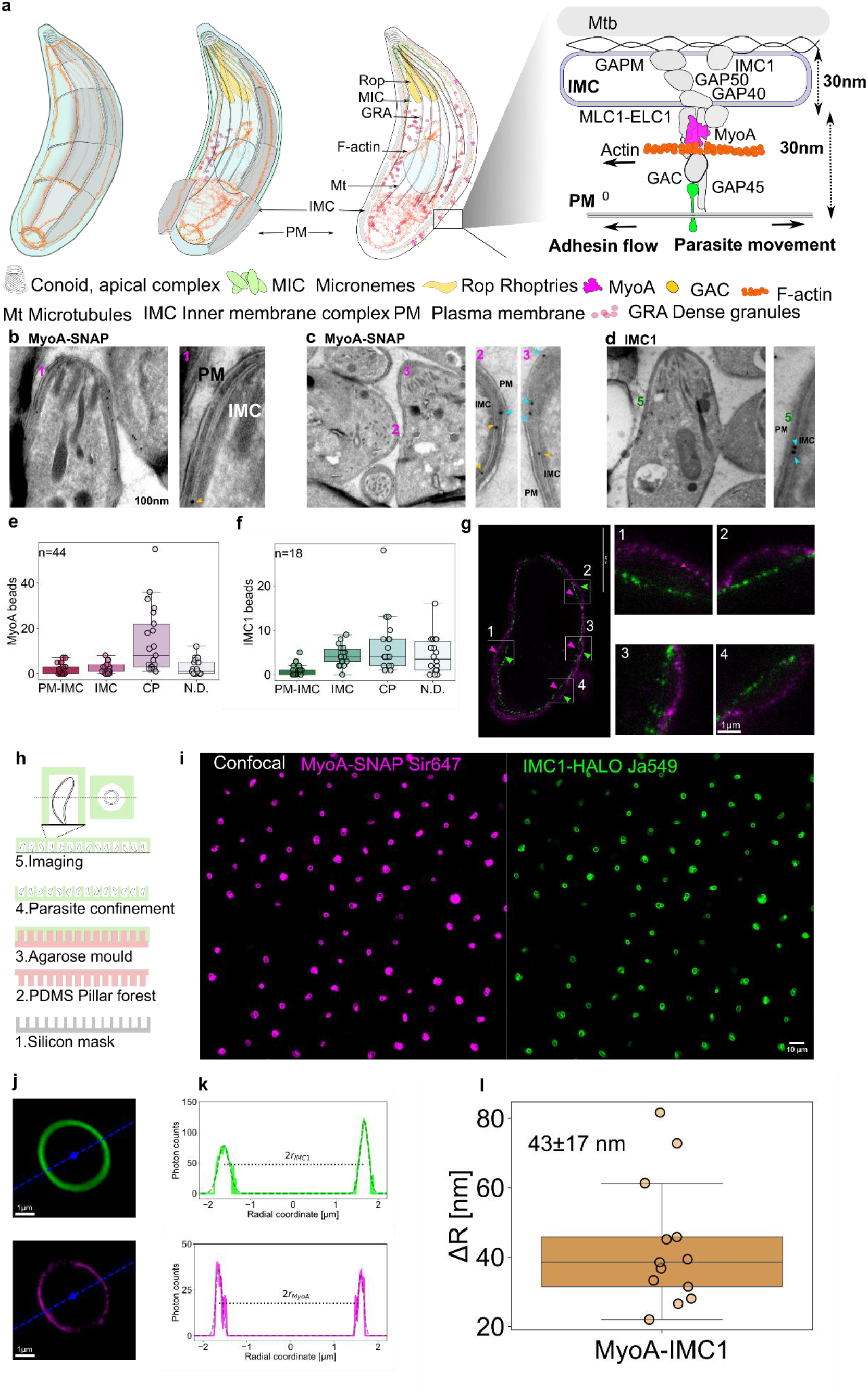
**a**. The linear motor model of migration proposes that the actin-myosin motor complex is localised in the space between the plasma membrane and the inner membrane complex (IMC), a structure consisting of membranous cisternae and structural components located ∼30 nm below the PM. Migration occurs when apical secretion of adhesion proteins (micronemes, rhoptries) and actin polymerisation from the apical end of the parasite engages with the myosin complex leading to parasite movement. Visualisation of MyoA in localized excentric to IMC1 labelled structures. **b-d**. Cryo immunolabeling according to Tokuyasu ^16^ of MyoA and IMC1 proteins labelled with 10 nm protein A colloidal gold supports a distribution of IMC1 populations in an inner localisation compared to MyoA. **e-f**. Quantitative distribution of MyoA, and IMC1. **g**. uExMIC of MyoA and IMC1 shows MyoA in an external position relative to IMC1. Microtubes for imaging with resolution Apicomplexa parasites. **h**. Recycled pillar forest confiners are used to create microtubes to vertically position individual parasites. **i**, Typical image of MyoA-SNAP IMC1-HALO tagged parasites in microtubes. **j**. For each cell, a centroid is estimated by fitting the fluorescent signal of the STED image with an ellipse. Then, the image is sectioned radially with multiple line profiles (only one is shown in the figure). **k**. For each line profile, a diameter is measured by fitting the fluorescent signal with two gaussians, one for each side of the cell. The distance between the two gaussians centers gives an estimate of the diameter. **l**. Estimated MyoA-IMC1 interdistance, computed as difference between the two respective radiuses estimated before, shows MyoA in an external position relative to IMC1. Mean and standard deviation are reported.

Migration occurs after the engagement of F-actin filaments with transmembrane proteins derived from secretion of micronemes (invasion related apical organelles) via the glideosome associated connector (GAC) to MyoA ^5^. The model has provided valid observations for translational approaches, but functional data still show a far more complex mechanism than suggested by the model with possible different motor configurations and actin-myosin independent mechanisms untested ^6^. Indeed, the orientation and localisation of the glideosome has in large part been inferred by immunoprecipitations ^7^, which can show direct interactions, but do not proof the topology of the glideosome. Therefore, the exact localisation of the motor is paradoxically not established, due to the limitation to resolve the motor complex with nanometric axial resolution.

In recent years, super-resolution and expansion microscopy methods have shattered the diffraction limit and resolution even below 20 nm in all dimensions can be achieved ^8 9 10^. However, many of the technologies have been demonstrated in proof of principle studies to test resolution in the third dimension but have not yet been used to localise the components of complex biological samples.

Here, we used a combination of super-resolution technologies, including Graphene energy transfer (GET) ^11^, uExMIC combined with STED microscopy in parasite confiners, and cryo-immunolabeling to determine the exact localisation of MyoA, the core motor of the glideosome. Our data present the first direct localization of Myosin A populations with nanometric resolution in the third dimension, which is at least partially compatible with the linear motor model.

## Results

Here we argue that we can put the linear motor model to its test by assessing whether a major claim of the model can be tested. Therefore, we implement complementary approaches, including GET, ^11^ to test whether the localisation of MyoA is, as predicted by the linear model, within the narrow space between the PM and IMC.

We visualise the localisation of MyoA with complementary approaches. Initially we use cryo-immunolabeling with 10 nm protein A colloidal gold. MyoA (Fig.1) and GAP45 (Fig.S1) gold particles localisations are found associated to the inter PM-IMC space, with an increased of IMC1 associated beads detected in the IMC. Next, we used ExSTED ^15^, a 4x expansion of tachyzoites in an hydrogel combined with STED. Expansion was not isotropic but made a distinct visible separation in x,y dimension for MyoA and the IMC1 structures (Fig. 1 and Fig. S1)

Next, we positioned the parasite vertically in casting micro-cylinders from pillar forest confiners that were repurposed for imaging, simplifying visualisation and quantification of pellicle markers to the lateral plane (for simplicity MIRA (microcilinders for imaging radially Apicomplexa parasites) and imaged with live STED (Fig. 1). The data provide an insight in the relative distribution of both proteins in live parasites. Since MIRA are repurposed pillar forest containers with dimensions bigger than the parasite, imaging was limited by the number of parasites positioned vertically in the confiner.

Width estimation using MyoA and IMC1 markers in both methods, show MyoA populations localised in the parasite outer side. Limitations of this approach was associated to the low density labelling of these markers and the conical geometry of the sample. Overall, results support that MyoA occupy an external position relative to IMC1 and the cytoskeleton microtubules (Fig. S1).

To precisely establish the relative position of MyoA and IMC1 we implement GET, that takes advantage of the optical properties of 2D graphene lattices, to measure z distances in the range of 40 nm above the coverslip down to ∼2 nm precision ^11^. Graphene is an universal energy transfer acceptor in the visible and near infrared spectrum behaving as a reversible quencher in a distance dependent manner. This property makes GET a precise nanometric ruler to test the localisation of labelled proteins in the PM-IMC region. We doubly-labelled MyoA and IMC1 proteins with SNAP and HALO respectively ^12 13^ using CRISPR-CAS9 ^14^ because they are defining markers for the motor and the IMC complex, the barrier that compartmentalize the PM-IMC space and the cytoplasm. Placing this cell line on graphene coated slides we measured changes in lifetime (Fig. 2) to calculate the averaged relative distance to the graphene of both markers (Fig. 2). The results show a MyoA population in the axial quenching range of graphene that is closer to the surface than IMC1 ^1^. The distance between MyoA and IMC1 is consistent between measurements (Fig. 2), and matches the IMC width measurements obtained with cryo-EM tomography, a method with higher resolution than light imaging but which does not discriminate between individual proteins. Overall, the results obtained with GET support that a population of MyoA and IMC1 maintain a constant distance to each other corresponding to fixed positions on outer and inner sides of the IMC respectively (Fig. 2).

**Figure 2.**
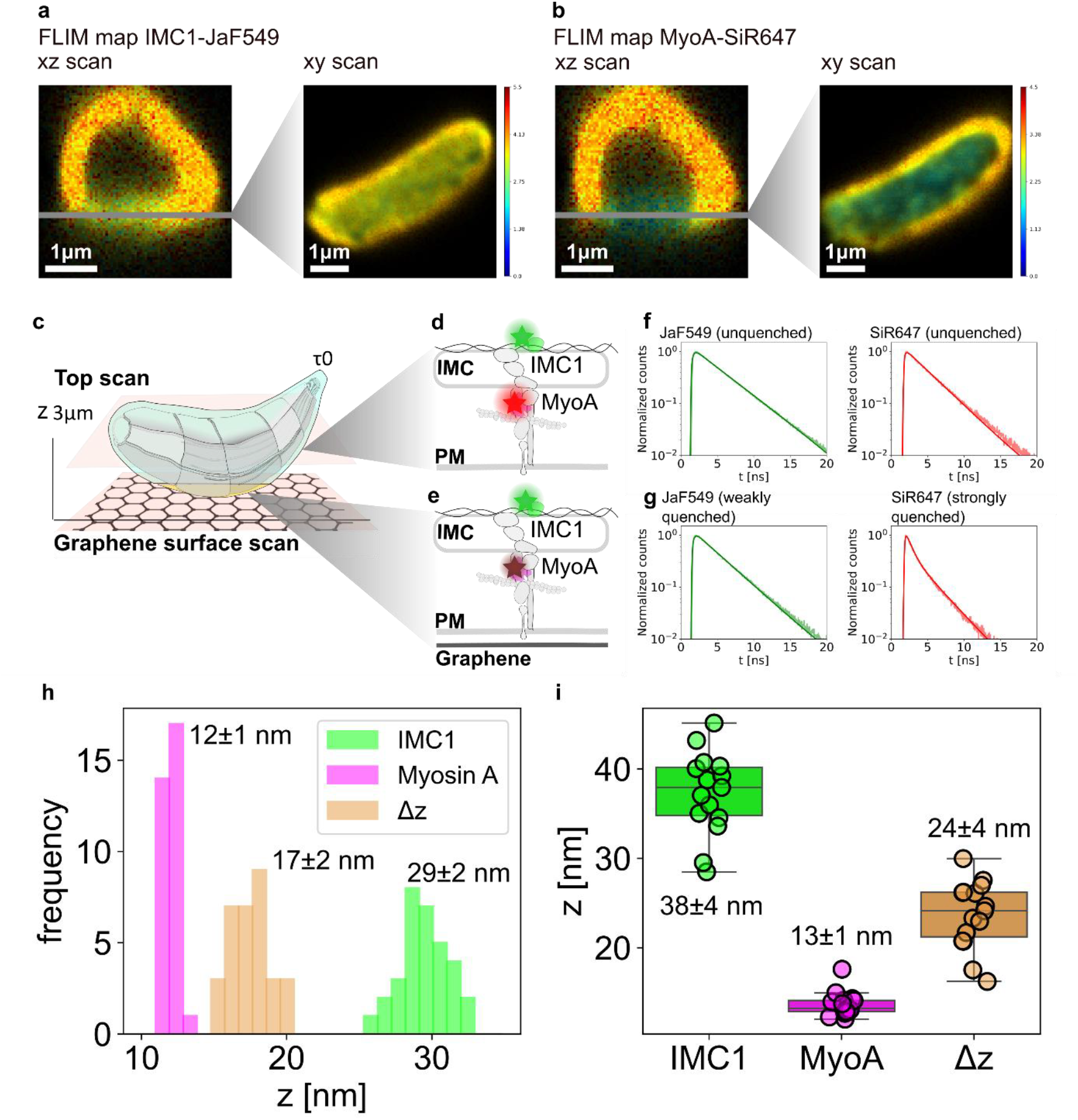
**a, b**. Vertical and horizontal FLIM maps of a doubly-labeled parasite. The vertical scan is used to identify the plane of graphene, where the horizontal scan is then taken. The colour-coded map shows a stronger quenching in the red channel on the graphene surface, and no quenching outside of the range. Note that the lifetime color-scale is adapted to the unquenched lifetime of the dye for better visual comparison, and hence differs in the two channels. **c**. Labeled parasites are positioned and fixated on a graphene substrate. The signal from the fluorescent markers is measured with a scanning confocal microscope both on the graphene surface and a few μm above it. **d, e**,. Labeled proteins in close proximity to graphene show varying degrees of fluorescent intensity and lifetime quenching depending on their relative distance to graphene, while proteins outside of the quenching range provide unquenched reference lifetime values. **e, f**. Example of fluorescent decay curves showing different degrees of (lifetime) quenching. Measurements beyond the quenching range of graphene in both channel show no quenching and are used as reference. Measurements on the graphene surface show very weak quenching in the green channel, suggesting an average z position close to the upper limit of the quenching range, and a strongly quenched component in the red channel, suggesting the presence of a Myosin population close to graphene. **h**. Histograms of protein-graphene and protein-protein relative distances for different surface points of the cell shown in **a, b. i**. Protein-graphene and protein-protein distances measured for several surface points of the same cell are then averaged; average values for all cells are then plotted together in **i**. The results support the presence of a myoA population in agreement with the linear motor model, with a distance to IMC1 consistent with localisation respectively on the outer and inner side of the IMC complex

## Discussion

Proof of principle experiments using DNA origami based nano-emitters show that novel GET is able to measure the z position with resolution up to ∼1 nm in a ∼40 nm Z range. This made GET an attractive tool to test directly whether the prediction of MyoA motor localisation made by the linear motor model was correct. GET results show a MyoA population positioned in the PM-IMC space with an axial localization precision of 1-4 nm, supporting the validity of the linear model.

Beside, taking MyoA and IMC1 as markers for the outer and inner side of the IMC, we use GET as a nanometric ruler to measure the z dimension of the IMC complex, something only amenable until now to EM based techniques, but with the advantage of protein specificity. GET and EM estimation of the IMC thickness show good agreement and comparable resolution. By comparison, with other super-resolution methods GET has superior z resolution than ExSTED, SIM, DNA-PAINT, or uExMIC without requiring the complex set up and sample optimisation of MinFlux.

The method has limited measurements yield. This is mainly due to heterogeneous parasite attachment and possible presence of debris, such as cell lysate, between the graphene surface and the cell membrane. These can increase the distance between the surface and the target proteins, which can fall outside of the quenching range. For these reasons, only perfectly attached cells could be considered in measurements. Further work will aim to investigate whether there are smaller subpopulations of the motor complex beyond the pellicle localisation implementing uExMIC cryofixation that minimising fixation artifacts in MIRA confiners designed de novo, rather than repurposing recycled pillar forests, with dimensions closer to the size and shape of Toxoplasma, following an scale up approach previously described in microfabrication devices for imaging bacteria ^17^.

## Acknowledgements

This study was supported by the Deutsche Forschungsgemeinschaft (DFG; German Research Foundation). Professor Olympia Ekaterini Psathaki, was supported by grants: German Research Foundation, DFG CRC 1557, Z-project, German Research Foundation DFG iBiOs PI405/14-German Research Foundation DFG Priority Program SPP2225 EXIT Strategies of Intracellular Pathogens. We thank Simon Gras for excellent technical assistance, Professor Jörg Renkawitz and Mr Mauricio Ruiz for assistance and sharing pillar forest confiners, and the Core Facility Bioimaging at the Faculty of Veterinary Medicine, the Core Facility Flow Cytometry, of the Chair of Experimental Parasitology, Faculty of Veterinary Medicine LMU for excellent support. G. F. thanks Dr. Alan Szalai for fruitful discussion.

## Author contributions

Giovanni Ferrari: Conceptualization; investigation; methodology; writing – review and editing.

Yuan Song: Investigation; review and editing.

Michael Hensel: Investigation; editing.

Katherina Pshataki: Investigation; methodology; review and editing.

Doan Thuy Nguyen Nguyen: Investigation; data acquisition.

Simon Gras: Generation of reagents.

Markus Meissner: Funding acquisition; review and editing.

Javier Periz: Conceptualization; supervision; investigation; writing – original draft, review, project administration and editing.

Philip Tinnefeld: Conceptualization; supervision; investigation; funding acquisition; project administration; writing – review and editing

## Supplementary Figure

**Figure. S1.**
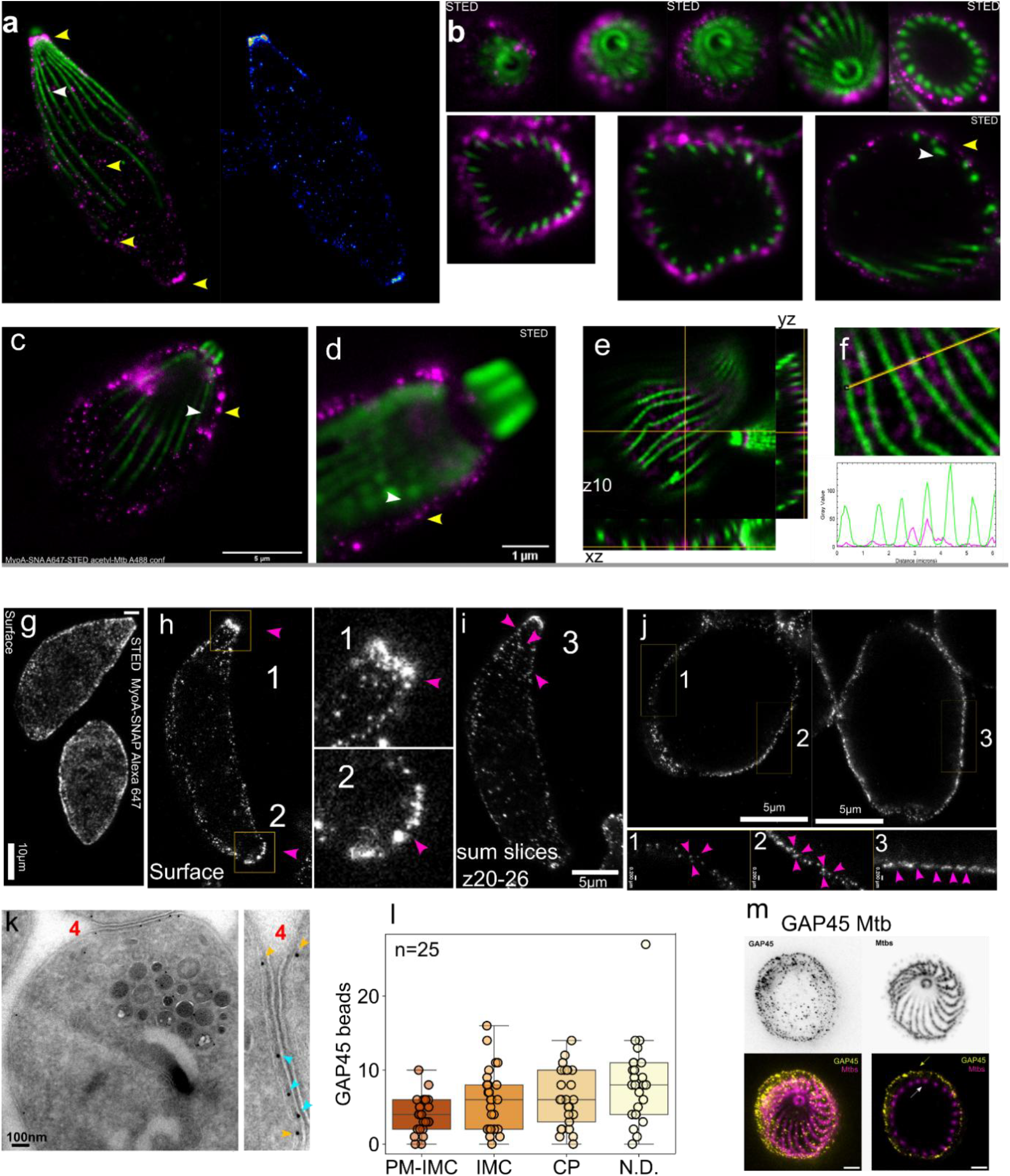
a) Distribution of MyoA (STED microscopy) and microtubules (confocal) in an expanded tachyzoite. b-c-d) Distribution of MyoA and microtubules in parasite sections (e-f) Plot distribution of MyoA and microtubules. (g-h-i-j). STED imaging of MyoA suggests a flexible distribution of the pellicle. (k-l) Cryo immunolabeling according to Tokuyasu of GAP45 proteins labelled with 10 nm protein A colloidal gold supports a distribution of GAP45 populations in the pellicle.

