## Supplementary material for "Localisation of Apicomplexa motor Myosin A with axial nanometric precision using graphene energy transfer": Materials and Methods

**Generation of transgenic parasites**

New strains were generated via the use of CRISPR/Cas9 as previously described ^4^. Guide RNAs targeting the regions of interest were designed using EuPaGDT ^9^. The sequences of all gRNAs and primers can be found in Supplementary Table 1. Briefly, gRNA oligos were annealed, ligated into the Cas9-YFP vector and verified by sequencing (Eurofins Genomics). Reparation templates DNA were generated by amplifying the Halo or YFP with 50 bp homology arms to the gene of interest by PCR using Q5 High-Fidelity DNA Polymerase (New England BioLabs). The repair template was purified using a PCR purification kit (Blirt, EM26.1). Tachyzoites were pooled with the repair template PCR and 10 μg of the corresponding Cas9 vector. Parasites were then transfected with Amaxa 4D-Nucleofector system (Lonza AAF-1003X). The transfected parasites were allowed to invade fresh HFFs and replicate for 48 hours. After manual egress and filtration through a 3μm filter, the parasites were enriched for those expressing Cas9-YFP using FACS (FACSARIA III, BD Biosciences) and sorted into 96-well plates. Proper integration of repair templates was confirmed via PCR and sequencing (Eurofins Genomics).

**Table 1**:

| IMC1 |  |
| --- | --- |
| gRNA forward | AAGTTGTACAAGTATACCACGCTGAG |
| gRNA reverse | AAAACTCAGCGTGGTATACTTGTACA |
| Homologies forward | GCGTCATGTGCGCGGTCTGCGGCGGTGATGGTGTGTGCCGATGCCAGTGC GCTAAAATTGGAAGTGGAGG |
| Homologies reverse | GAGGTCAGATTCCAGAAATTCCCGTCGCACAAGGCCTTCAGCGTGGTATA ATAACTTCGTATAATGTATGCTATACG |
| Integration forward | GATCGAAGAGGCACATGCTG |
| Integration reverse | ACTTGAAATTCCCGTGCCAG |
| MyoA |  |
| gRNA forward | AAGTTAACGAACGTGTCTAGAACGCG |
| gRNA reverse | AAAACGCGTTCTAGACACGTTCGTTA |
| Homologies forward | gACACCTGGTGGACAACAACGTCAGCCCCGCGACTGTTCAGCCGGCGTTCGCTAAAATTGGAAGTGGAGG |
| Homologies reverse | AGAGAAAACAACAAATGGATGAGAAGAGTTCAAAAGGCAAACGAACGTGTATAACTTCGTATAATGTATGCTATACG |
| Integration forward | cttggccgttgtttctcc |
| Integration reverse | ACAAAGAAGCGCACAAACG |

**MIRA (microcilinders for imaging radially Apicomplexa )**

PDMS Pillar forest confiners for studying cell navigation were a gift from Professor Renkawitz lab, and the preparation and design has already been published ^1^ Briefly, Polydimethylsiloxane (PDMS) was mixed, elastomer and curing agent 1:10, poured over silicon wafer with the negative microchannel structures imprinted by photolithography, degassed in vacuum, and cured at 80 °C overnight. After that, the PDMS mould was then used as a second positive wafer to cast agarose micro-cilinders for MIRA.

Agarose mould were cast with 2% Agarose molecular biology grade dissolved in 1xTAE using the generated PDMS pillar forest confiners of 5 by 12 μm and 12 microns distance. Labelled parasites resuspended in Fluorobrite media at 1.10^8^.ml^-1^ added 50 μl to the MIRA devices at a concentration , spinned for 3000 rpm for 5 minutes, the excess of media removed with a tissue and the gel mounted on glass (20mm dish ♯ 1.5; 0.16-0.19 mm, Cellvis) that has been previously coated with d-polylysine, left dried and wash 3x with PBS. Imaging was made with an 3D confocal-STED Abberior.

**Image Analysis**

Fiji were used for image processing. All Myoa-IMC1 experiments were performed three independent biological replicates and repeated a fourth time with secondary antibodies labelled with dyes emitting in the opposite emission spectrum . Except cryo-immunolabelling Electron microscopy experiments for IMC1 and GAP45 (one experiment and two experiments respectively). Statistical analysis was conducted using GraphPad Prism (version 8.0.1) using tests as stated in the figure legend.

**Tagging of parasites**

Toxoplasma cell line parasite line ΔQ80m strain was tagged with HALO ^2^ and SNAP ^3^ following standard protocols established in the lab ^4^. Parasites were maintained on HFFs (Human foreskin fibroblasts (HFFs) (RRID: CVCL_3285, ATCC) in DMEM ( Dulbecco's modified Eagle's medium), 10% foetal bovine serum, 2 mM l‐glutamine and 25 mg.ml^-1^ gentamicin, and maintained at 37°C and 5% CO2. Cultured cells and parasites were screened against mycoplasma contamination with LookOut Mycoplasma detection kit (Sigma) and Mycoplasma Removal Agent (Bio‐Rad) if needed.

**8. Parasite staining for MIRA and graphene assays**

HFFs 4cm dish cell cultures were infected with 110^8^ parasites and left to replicate for 48h until replicating vacuoles were ready to egress. Parasites expressing MyoA-SNAP IMC1‐HALO were incubated sequentially with Janelia dye 549 HALO ligand (Promega) and Sir647 SNAP ligand (NEB) at a final concentration of 20 nM and 500 nM respectively for 1h. Wash 3x in complete DMEM media for 30 min.

After that parasites were mechanically released scratched, syringed , filtered following established protocols in the lab ^2^, parasites were centrifuged at 300g for 5 min and resuspended in PBS 37°C for an additional washing step. After that the parasites were spinned down again resuspended in warmed PBS media, at approximately 1.10^8^.ml^-1^ and added 50μl to a graphene slide for 30 min, then removed carefully, check for attachment to the surface and fixed ^5^ or alternatively used in confinement experiments with MIRA devices.

Imaging	was done with a 3D STED Abberior microscope. To measure MyoA and IMC1 we use 2D ExSTED with optimised setting powers for these dyes, as a guide (10% laser power), x,y 15-15 nm pixel size , dwell time 5 μs, 7 line accumulations.

**9. uExMIC** for uExMIC parasites were processed following published protocols ^6^ with the following differences MyoA and IMC1 proteins were labelled using primary antibodies rabbit polyclonal anti SNAP 5μg.ml^-1^ (Biozol) and mouse monoclonal anti IMC1 dilution 1:200 (a gift from Professor Gary Ward, University of Vermont), acetylated microtubule monoclonal mouse antibody 2μg.ml^-1^ (Sigma-Aldrich) from and using as a secondary antibodies donkey anti rabbit Alexa 647, donkey anti mouse Alexa 594 and Alexa 488 from Invitrogen at 4 μg.ml^-1^. Expansion was repeated a fourth time using as secondary antibodies anti rabbit goat Abberior 580 and goat anti mouse Abberior 635P yielding reproducible MyoA-IMC1 ratio results.

To measure MyoA and IMC1 we use 2D STED ( 3D STED Abberior microscope) mode with optimised setting powers for these dyes, as a guide (1% laser power), 40-40 nm pixel size , dwell time 5 μs, 10 line accumulations, with an 100 objective 1.4 NA, pin hole 1A.U, immersion oil Zeiss F30cc, type F.

MyoA IMC1 measurements with uExMIC were done in four independent experiments with consistent results, 14 images measured. In case of having sufficient separation between between MyoA and IMC1, we use confocal mode for estimating the width ratio of the parasites confocal mode ( 3D STED Abberior microscope) with optimised setting powers for these dyes, as a guide (2% and 15% laser power to image dye Abberior 580 and 635Prespectively ), 60-60 nm pixel size , dwell time 5 μs, 5 line accumulations . To measure MyoA and Mtb we use a STED and confocal mode respectively. With power conditions optimised for each image. Three independent experiments were done and more than 100 images taken.To estimate the relative position of MyoA and IMC1 we took five width measurements in the apical, mid and posterior end and calculate a relative ratio, >1< myoA width above or below reference marker IMC1.

**Cryo sample preparation for Tokuyasu** ^7^ ^8^ **immunogold labeling and TEM**

Cells were fixed for 2 hours at RT in 2% (w/v) formaldehyde/0.1% (v/v) glutaraldehyde in 0.1M PB buffer pH 7.4. For embedding, cells were washed with 0.1% glycin in 0.1M PB buffer pH 7.4 followed by 0.1M PB buffer pH 7.4 and then centrifuged (300g, 3 min) in 1% (w/v) gelatin/0.1M PB buffer pH 7.4. The pellet was infiltrated at 37°C in 10% (w/v) gelatin/0.1M PB buffer pH 7.4. After gelation of the gelatin-embedded sample on ice, 1 mm3 sample cubes were dissected and transferred to 2.3 M sucrose in 0.1M PB buffer pH 7.4 for infiltration overnight at 4°C. The cubes were mounted on aluminium stubs, plunge frozen in liquid nitrogen and then placed in the cryo-ultramicrotome (UC7, Leica Microsystems, Wetzlar). Ultrathin 60 nm sections were cut at -110°C using a cryo-immuno diamond knife (Diatome, Switzerland). Ribbons of cryosections were collected in a Perfect Loop (Science Services, Germany) with a 1:1 mixture of 1% (w/v) methylcellulose and 2.3M sucrose in 0.1M PB buffer pH 7.4 and transferred to Formvar carbon-coated 50µm mesh copper grids.

For immunolabelling, grids containing sections were placed on 37°C pre-warmed Milli-Q water for 45 minutes to melt the methylcellulose/sucrose solution and gelatin. Grids with sections were rinsed 8 times with 0.1% glycine in PBS pH 7.4 (8 times), blocked for 3 min in 1% BSA in PBS pH 7.4, incubated for 60 min in primary antibodies: rabbit anti SNAP from Biozol, rabbit anti GAP45 ( a gift from Professor Dominique Soldati -Favre, University of Geneva), or mouse anti IMC1 ( a gift from Professor Gary Ward, University of Vermont ) diluted in 1% BSA, 0.2% fish skin gelatin in PBS pH 7.4, washed 8 times in 0.1% BSA in PBS pH 7. 4 and then incubated for 30 min with rabbit anti-mouse (BioZol original from Jackson-Immuno-315-005-048) as a bridging antibody (only necessary if the first Ab is not rabbit), diluted in 1% BSA, 0.2% fish skin gelatine in PBS pH 7.4. The grids were then washed 8 times in 0.1% BSA in PBS pH 7.4 and incubated in Protein A gold-10nm (CMC-Utrecht, batch 08-2021) in 1% BSA in PBS pH 7.4 for 20 min and fixed in 1% (v/v) glutaraldehyde in PBS pH 7.4 for 5 min, washed and stained 5 min on a drop of 2% uranyl oxalate (pH 7.0), followed by 10 min in 1,8% (v/w) methyl cellulose/0.4% uranyl acetate (pH 4.0) mixed 1:1 on ice. Sections were analyzed in a transmission electron microscope JEM 2100Plus at 200kV (JEOL, Japan) equipped with a XAROSA CMOS 20 Megapixel Camera (Emsis GmbH, Germany) and a Zeiss TEM 902 at 80kV.

**Graphene-on-glass coverslip preparation**

Monolayer graphene on a 60 mm × 40 mm copper substrate with poly(methyl methacrylate) (PMMA) on top was purchased from ACS Materials LLC, USA or Graphenea Inc., Spain. The foil was stored in a vacuum-sealed desiccator. A wet-transfer approach was used to transfer the graphene to glass coverslips (Paul Marienfeld GmbH & Co. KG, Germany).^21^ All coverslips were treated with ultrasound at 37˚C for three times 15 minutes in a bath of Milli-Q water to remove any contaminants from the surface. In the first of the three rounds, 1% v/v of Hellmanex^®^ III (Hellma GmbH & Co. KG, Germany) was added to the bath.

Smaller pieces of roughly 0.25 cm² were carefully cut from the PMMA/graphene/copper foil. The copper was wet etched by letting a piece float with the copper film exposed to 0.2 M ammonium persulfate ((NH_4_)_2_S_2_O_8_) for ~4 h. A coverslip was dipped vertically while slowly moving towards the PMMA/graphene, scooped gently out of the solution, and transferred to Milli-Q water in order to wash out the residues of ammonium persulfate. The step of washing PMMA/graphene was repeated twice with fresh Milli-Q water. Next, the PMMA/graphene was scooped with a glass coverslip, carefully dried using a nitrogen stream and stored overnight. Extra PMMA (M_w_ = 120,000 g/mol) was dissolved in chlorobenzene (50 mg/mL) and was drop-casted on top of the first PMMA/graphene layer. This allowed the dried PMMA to re-dissolve, thus relaxing the underlying graphene monolayer and forming an improved contact with the substrate ^64^. The added PMMA dried for at least 30 minutes.

For cleaning, the PMMA/graphene on glass was first dipped in two distinct acetone baths for 7 min each. Subsequently, the washing was repeated in toluene for 7 min. After each washing step, the samples were dried with a nitrogen stream. Finally, the graphene-on-glass was placed on active coal that was heated on a heating plate to 230 ˚C. Here, the graphene faces the active coal. The final heating was done for about 10 h prior to usage.

After this step, the graphene substrate was ready to be used. A chamber (SecureSeal™ hybridization chambers, Grace Bio-Labs, USA) was glued around the sheet of graphene on the glass coverslip and the cells were incubated and fixated as described in the corresponding section.

### **Confocal Fluorescence Microscope**

All experiments involving graphene were performed on a home-built confocal microscope based on an Olympus IX-71 inverted microscope. Here, the sample was excited by pulsed interleaved excitation with a 532 nm and a 636 nm lasers (LDH-P-FA-530B and LDH-D-C-640; both PicoQuant GmbH, Germany) at a frequency of 20 MHz (PDL 828 “Sepia II”, PicoQuant GmbH with an oscillator module: SOM 828, PicoQuant GmbH, Germany). A delay of 25 ns was set between the two laser pulses, and each channel detection was timegated with a 25 ns window. The lasers were coupled into a single mode fiber (P3-488PM-FC, Thorlabs GmbH, Germany) to obtain a Gaussian beam profile and to perfectly overlay the two excitation beams. Circular polarized light was obtained by a linear polarizer (LPVISE100-A, Thorlabs GmbH, Germany) and a quarter-wave plate (AQWP05M- 600, Thorlabs GmbH). The light was focused on a diffraction-limited spot using an oil-immersion objective (UPLSAPO100XO, NA 1.40, Olympus Deutschland GmbH, Germany). The position of the sample is adjusted using a piezo stage (P‑517.3CD, Physik Instrumente (PI) GmbH & Co. KG, Germany) and a controller (E-727.3CDA, Physik Instrumente (PI) GmbH & Co. KG, Germany). The emission light was separated from the excitation beam by a dichroic beamsplitter (zt532/640rpc, Chroma Technology Corporation, USA) and focused onto a 50 μm diameter pinhole (Thorlabs GmbH, Germany). After the pinhole, signals of different wavelengths were separated by a dichroic beamsplitter (640 LPXR, Chroma Technology Corporation, USA) into a green (Brightline HC582/75, AHF, Germany; RazorEdge LP 532, IDEX Health & Science, LLC, USA) and red (SP 750, AHF, Germany; RazorEdge LP 647, IDEX Health & Science, LLC, USA) detection channel. The emission was focused onto avalanche photodiodes (SPCM-AQRH-14-TR, Excelitas Technologies Corporation, USA) and the signals were registered by a time-correlated single photon counting (TCSPC) unit (HydraHarp400, PicoQuant GmbH, Germany). The setup was controlled by a commercial software package (SymPhoTime64, Picoquant GmbH, Germany). The laser power was adjusted for each color accordingly to the sample labeling density but always in the 0.2-0.5 μW range.

### **Fluorescence Lifetime Measurements**

FLIM maps of cells were obtained by scanning the sample with the piezo stage with a fixed pixel size of 50 nm and a dwell time per pixel of 2 ms. The size of the scanned area was adjusted according to the size of the cell. The local fluorescence lifetime is estimated pixel by pixel through the median value of the detected photon arrival times. Finally, the local fluorescence lifetime is color coded and a FLIM map of the cell is obtained.

Once a relevant cell was identified on the graphene coated coverlip, a vertical (XZ or YZ) scan was acquired to locate the graphene plane. Then, on that plane, multiple (10-40) spots were chosen at a distance of 4-500 nm covering the whole bottom membrane of the cell. Each spot corresponded to a 5 s confocal measurement, during which photons were collected. Later, a single fluorescence decay fit was performed for each one of these measurement, yielding an estimation for the protein z position in that spot.
